# SynOmics: Integrating Multi-omics Data Through Feature Interaction Networks

**DOI:** 10.1101/2025.09.17.676954

**Authors:** Muhtasim Noor Alif, Wei Zhang

## Abstract

The integration of multi-omics data is essential for achieving a comprehensive understanding of molecular systems and enhancing the performance of predictive models in biomedical research. However, many existing models have limited capacity to capture cross-omics feature interactions, which hinders the depth of integration. In this study, we introduce SynOmics, a graph convolutional network framework designed to improve multi-omics integration by constructing omics networks in the feature space and modeling both within- and cross-omics dependencies. By incorporating both omics-specific networks and cross-omics bipartite networks, SynOmics enables simultaneous learning of intra-omics and interomics relationships. Unlike traditional approaches that rely on early or late integration strategies, SynOmics adopts a parallel learning strategy to process feature-level interactions at each layer of the model. Experimental results demonstrate that SynOmics consistently outperforms state-of-the-art multi-omics integration methods across a range of biomedical classification tasks, highlighting its potential for biomarker discovery and clinical applications.

**Availability:** Source code is available at https://github.com/compbiolabucf/SynOmics.

**Supplementary information:** Supplementary data are available at *Journal Name* online.

## Introduction

Biological systems and diseases are influenced by complex interactions among various omics types. To gain a deeper understanding of complex diseases, it is crucial to comprehend how these omics types interact both within and across their respective layers [1, 2]. Multi-omics integration is essential for effectively capturing these interactions [3, 4]. With the advancement of high-throughput sequencing technologies, researchers can now generate high-quality, precise multi-omics data [5]. While each omics technology provides only a partial view of biological complexity, integrating multiple omics types offers a more comprehensive understanding of the underlying biological processes. However, this integration remains a challenging task due to the complex relationships among the different omics types.

Over time, a wide range of methods have been developed for integrating multi-omics data, enabling various downstream applications such as disease and cancer research, biomarker discovery, and drug development [3, 4, 6, 7, 8]. These approaches leverage complementary information from multiple omics types—such as genomics, transcriptomics, proteomics, and metabolomics—to provide deeper biological insights and improve the accuracy of predictions in biomedical research.

Since molecular structures and interactions often exhibit graph-like patterns [9, 10, 11], the rise of graph-based deep learning techniques has led to the widespread use of graph neural networks (GNNs) [12] for multi-omics data integration [13, 14,15].

Wang et al. [16] introduced the Sample Similarity Network Fusion (SNF) approach to create a unified graph representation across multiple omics types. Although SNF does not involve deep learning, it has been widely adopted to generate joint networks that can serve as inputs to GNN models. Building on this, Li et al. [17] proposed MoGCN, which extends SNF by incorporating a graph convolutional network (GCN) [18] for predictive modeling. MoGCN employs an early integration strategy using autoencoders to directly combine input omics layers and learn a joint representation. However, this approach does not separately model omics-specific relationships, potentially overlooking important intra-omics information. In contrast, late integration methods aim to capture omics-specific interactions before aggregating information across omics types. For example, MOGONET [19] uses omics-specific GCNs to learn intra-omics representations and then integrates these embeddings through a correlation network. Similarly, SUPREME [20] trains omics-specific GCNs independently before merging the resulting embeddings for final prediction. While these approaches typically rely on fixed sample similarity networks, MOGLAM [21] employs a dynamic graph convolutional approach that updates the similarity networks as the model learns omics-specific interactions.

Most existing models construct sample similarity networks to facilitate learning by sharing information across samples [17, 19, 20, 22, 23]. While this approach can be effective, it has several notable limitations. One major drawback is its inability to capture the inherent interactions among omics features and molecular entities—interactions that exist independently of individual samples and are fundamental to biological systems. Although sample similarity networks may reflect broad population-level patterns, they often overlook critical feature-level mechanisms such as gene regulation and pathway dependencies, resulting in the loss of important biological insights. Furthermore, omics data are typically characterized by high dimensionality and a limited number of samples. This imbalance, where the number of features greatly exceeds the number of samples, poses a significant challenge for effective integration. In such cases, sample similarity networks often fail to capture the complex dynamics and relationships that underlie real-world biological systems.

Another limitation of traditional approaches is their tendency to model only intra-omics relationships using graph-based learning, while treating inter-omics dependencies separately through domain-agnostic architectures such as multilayer perceptrons (MLPs) or standard autoencoders [17, 19, 20, 23]. This decoupled design restricts the model’s ability to capture coordinated signals across omics types, ultimately limiting the quality of the learned representations. In biological systems, omics layers are tightly interconnected—genomic alterations can drive transcriptomic changes, which may in turn influence epigenetic modifications or proteomic responses. Ignoring these interdependencies or modeling them in a disjointed manner can result in fragmented or biologically shallow representations. GNNs offer a promising alternative by enabling structured information flow across omics layers. However, their application in cross-omics integration remains limited, primarily due to the challenge of defining biologically meaningful interaction networks between heterogeneous omics types—for example, linking mRNA targets to their regulatory microRNAs (miRNAs). While intra-omics networks often leverage well-established resources such as functional interaction databases or co-expression networks, cross-omics relationships are typically less welldefined. Constructing these inter-omics connections often requires integrating prior biological knowledge, curated interaction databases (e.g., miRNA-mRNA targeting), or statistical inference from multi-omics datasets. In the absence of robust strategies for defining these cross-layer links, the full potential of GNNs for cross-omics integration remains underutilized.

In this study, we introduce SynOmics, a GCN-based framework designed to enhance multi-omics integration through feature-level learning. Our method integrates both intra-omics and inter-omics information by leveraging omics-specific networks and cross-omics bipartite networks. By operating in the feature space rather than the sample space, SynOmics provides a more nuanced representation of biological interactions. The bipartite networks offer a structured approach for capturing inter-omics information flow [24, 25, 26, 27]. Unlike conventional early or late integration strategies, SynOmics employs a parallel learning approach to jointly model within- and cross-omics patterns. Experimental results demonstrate that SynOmics performs robustly across a variety of tasks, highlighting its potential for heterogeneous data integration, clinical biomarker discovery, and improved clinical diagnostics.

## Method

In this section, we first describe the standard formulations of graph convolutional and bipartite graph convolutional layers. We then present SynOmics, our proposed framework for robust multi-omics integration, which leverages feature-level graph convolution both within and across omics types. Finally, we outline the evaluation methods used to assess the performance of SynOmics.

### Graph Convolution

Omics data often exhibit graph-like structures due to complex regulatory mechanisms, making graph convolutional networks (GCNs) [18] well-suited for modeling these interactions through efficient node embedding and message propagation. Accordingly, SynOmics employs graph convolution to capture intra-omics interactions within each omics type.

In traditional GCNs, 𝒢 = (𝒱, ℰ, 𝒳) is an attributed graph, where 𝒱 = {*v*_1_, *v*_2_, …, *v*_*n*_} is the set of nodes, *E* is the set of edges between the nodes, and **X** = {*x*_1_, *x*_2_, …, *x*_*n*_} ∈ ℝ^*n×d*^ is the feature matrix, where each row *x*_*i*_ represents the feature vector of node *v*_*i*_. The adjacency matrix for an unweighted graph is defined as **A** ∈ {0, 1}^*n×n*^, where *A*_*ij*_ = 1 if there is an edge between node *v*_*i*_ and node *v*_*j*_ and *A*_*ij*_ = 0 otherwise.

Then the *l*^*th*^ layer operations are specified by:

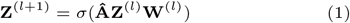

where **Z**^(*l*+1)^ is the output feature matrix of the *l*^*th*^ layer, *σ*(·) is the non-linear activation function, **Â** is the normalized version of the adjacency matrix **A, W**^(*l*)^ is the associated weight matrix of layer *l*, **Z**^(*l*)^ is the input feature matrix and notably, **Z**^(0)^ = **X**. The normalized adjacency matrix **Â** is defined by:

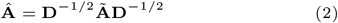

where **Ã** = **A** + **I** denotes the adjacency matrix with self connections, and **D** is the degree matrix of **A**.

### Bipartite Graph Convolution

To account for the influence of one omics type on another, we extend GCNs to bipartite networks for modeling cross-omics interactions. This approach, known as bipartite graph convolution (BGCN), has been adopted in prior studies [28, 29, 30, 31] and enables structured information flow between two distinct datasets.

Let 𝒰 and 𝒱 be two sets of nodes, where 𝒰 = {*u*_1_, *u*_2_, …, *u*_*n*_} and 𝒱 = {*v*_1_, *v*_2_, …, *v*_*m*_}. Then, the bipartite graph is defined as ℬ = (𝒰, 𝒱, ℰ_ℬ_), where ℰ_ℬ_ ∈ 𝒰 *×* 𝒱 is the set of edges between *U* and *V*. Let **B**_*u*_ ∈ {0, 1}^*n × m*^ the incidence matrix from 𝒰 to 𝒱 and **B**_*v*_ ∈ {0, 1}^*n × m*^ the incidence matrix from 𝒱 to 𝒰, which is notably the transpose of **B**_*u*_ for an undirected graph. 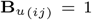 if there is an edge between nodes *u*_*i*_ and *v*_*j*_, and 0 otherwise. We define the adjacency matrix of the bipartite graph **B** ∈ {0, 1}^(*n*+*m*)*×*(*n*+*m*)^ as:

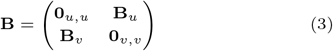

where **0**_*u,u*_ ∈ {0}^*n×n*^ and **0**_*v,v*_ ∈ {0}^*m×m*^ are zero matrices. Given the incidence matrices, we define the *l*^*th*^ layer of bipartite graph convolution as following:

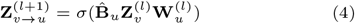

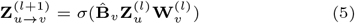

where, 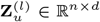 and 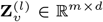 are the feature matrices of 𝒰 and 𝒱, respectively. 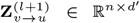 and 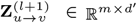 are the hidden representations of nodes in 𝒰 and 𝒱, aggregated from the opposite domain. *σ*(.) denotes the non-linear activation function. 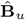 and 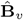 are the normalized incidence matrices of **B**_*u*_ and **B**_*v*_. 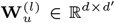 and 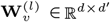 are the associated weight matrices. The adjacency matrix **B** in equation (3) is normalized in the same way as equation (2), and then the incidence matrices are extracted to get the normalized incidence matrices 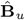 and 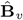.

### SynOmics

SynOmics is a supervised framework for multi-omics integration in biomedical classification tasks (Fig.1). It employs graph convolution for intra-omics learning and bipartite graph convolution for modeling inter-omics regulatory interactions. Unlike approaches that rely on sample-level similarities, SynOmics focuses on biologically meaningful regulatory links between features, such as miRNA regulation of mRNA expression, to capture more relevant signals. The framework operates on feature-level networks, where nodes represent molecular features and edges represent their biological relationships. To support this, we adapt the standard GCN and bipartite GCN formulations accordingly. The mathematical notations used in SynOmics are summarized in Table 1.

**Fig 1.**
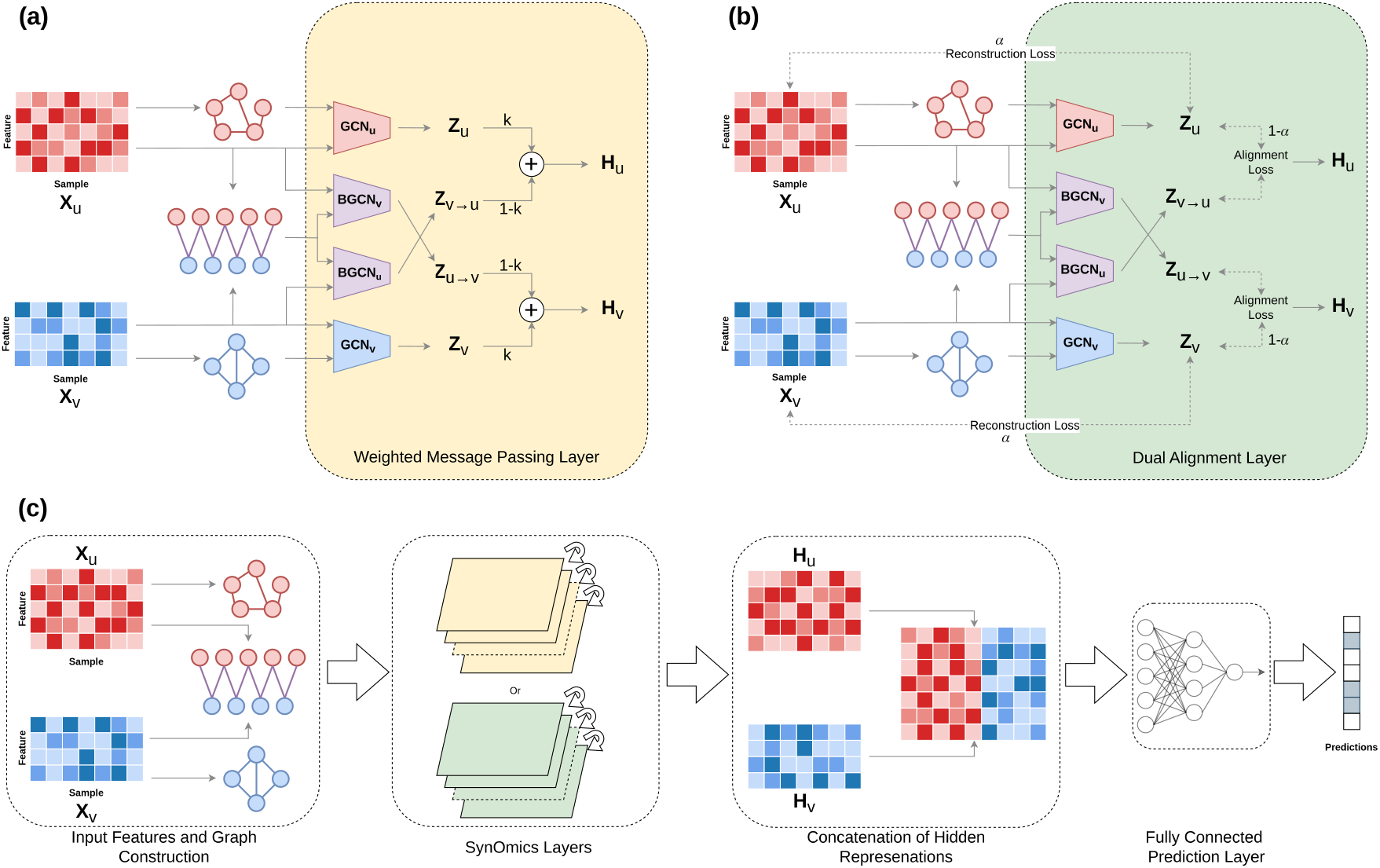
Illustration of SynOmics. SynOmics integrates two or more omics datasets by leveraging omics-specific feature networks and inter-omics bipartite networks. At each layer, the model applies two intra-omics GCNs and two inter-omics bipartite GCNs for each omics pair, using both the feature matrices and the corresponding networks. For each omics dataset, SynOmics generates two distinct embeddings: one capturing intra-omics representations and another modeling inter-omics relationships. (a) The weighted message passing layer combines the intra-omics and inter-omics embeddings through weighted aggregation to produce the layer output. (b) The dual alignment layer aligns the intra-omics and inter-omics embeddings with each other and with the original input features to produce the output. (c) The overall SynOmics architecture concatenates the final-layer outputs from either (a) or (b), followed by a fully connected module to generate the final prediction. Note: For illustrative clarity, matrices are presented in reduced form; in practice, omics feature spaces are considerably higher in dimensionality. *Alt Text: SynOmics diagram illustrating intra- and interomics GCN modules and two downstream strategies—weighted message passing and dual alignment—for integrating learned representations and producing final predictions*.

**Table 1.** Mathematical notations for SynOmics.

| Name | Definition |
| --- | --- |
| $n$ | number of samples |
| $p$ | number of features of omics1 |
| $q$ | number of features of omics2 |
| $nc$ | number of classes |
| $\mathbf{X}_u \in \mathbb{R}^{n \times p}$ | feature matrix of omics1 |
| $\mathbf{X}_v \in \mathbb{R}^{n \times q}$ | feature matrix of omics2 |
| $\mathbf{A}_u \in \{0, 1\}^{p \times p}$ | adjacency matrix for omics1 features |
| $\mathbf{A}_v \in \{0, 1\}^{q \times q}$ | adjacency matrix for omics2 features |
| $\mathbf{B}_u \in \{0, 1\}^{p \times q}$ | bipartite incidence matrix from omics1 to omics2 |
| $\mathbf{B}_v \in \{0, 1\}^{q \times p}$ | bipartite incidence matrix from omics2 to omics1 |
| $\mathbf{W}_u^{(l)} \in \mathbb{R}^{p \times p}$ | weight matrix of GCN<sub>u</sub> of the $l^{th}$ layer for omics1 |
| $\mathbf{W}_v^{(l)} \in \mathbb{R}^{q \times q}$ | weight matrix of GCN<sub>v</sub> of the $l^{th}$ layer for omics2 |
| $\mathbf{W}_{v \rightarrow u}^{(l)} \in \mathbb{R}^{p \times p}$ | weight matrix of BGCN<sub>u</sub> of the $l^{th}$ layer from omics2 to omics1 |
| $\mathbf{W}_{u \rightarrow v}^{(l)} \in \mathbb{R}^{q \times q}$ | weight matrix of BGCN<sub>v</sub> of the $l^{th}$ layer from omics1 to omics2 |
| $\mathbf{Z}_u^{(l+1)} \in \mathbb{R}^{p \times n}$ | intra-omics output feature matrix of the $l^{th}$ layer for omics1 |
| $\mathbf{Z}_v^{(l+1)} \in \mathbb{R}^{q \times n}$ | intra-omics output feature matrix of the $l^{th}$ layer for omics2 |
| $\mathbf{Z}_{u \rightarrow v}^{(l+1)} \in \mathbb{R}^{q \times n}$ | inter-omics output feature matrix of the $l^{th}$ layer from omics1 to omics2 |
| $\mathbf{Z}_{v \rightarrow u}^{(l+1)} \in \mathbb{R}^{p \times n}$ | inter-omics output feature matrix of the $l^{th}$ layer from omics2 to omics1 |
| $\mathbf{H}_u^{(l+1)} \in \mathbb{R}^{p \times n}$ | aggregated output feature matrix of the $l^{th}$ layer for omics1 |
| $\mathbf{H}_v^{(l+1)} \in \mathbb{R}^{q \times n}$ | aggregated output feature matrix of the $l^{th}$ layer for omics2 |

With the updated mathematical notations, we define the intra-omics graph convolution as:

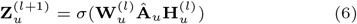

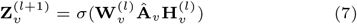

It is important to note that 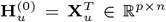 and 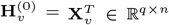 are the inputs for the first layer. Equations (6) and (7) enable information propagation by aggregating feature information from each node in one omics layer to its connected neighbors, including itself through self-connections.

The inter-omics bipartite graph convolution is defined as:

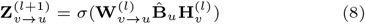

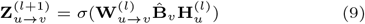

Equations (8) and (9) enable cross-omics information propagation from nodes in one omics layer to their connected counterparts in the other omics layer, as defined by the bipartite network. Specifically, in equation (8), 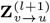 aggregates information for omics1 by collecting signals from 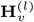, the hidden representation of omics2 from the previous layer, using the normalized bipartite incidence matrix 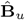. We apply the Leaky ReLU activation function [32] to obtain hidden representations. SynOmics executes the operations defined in equations (6) through (9) within a single layer, enabling parallel learning of intra- and inter-omics interactions as an alternative to staged integration strategies.

### Training

To investigate how training methodology influences downstream performance, we explore two alternative strategies for training SynOmics. The first approach, ‘weighted message passing’, balances the contributions of the intra- and inter-omics modules, providing a straightforward end-to-end training mechanism. This strategy is computationally efficient, faster to train, and easily scalable, making it ideal for rapid experimentation. The second approach, ‘dual alignment’, involves pretraining the GCN layers to enable the model to learn more detailed and comprehensive feature representations. By fine-tuning omics features before applying them to downstream tasks, this method enhances generalization and robustness. The details of these two training strategies are described in the following subsections.

### Weighted Message Passing

For effective integration, it is important to recognize that the two omics signals being considered may not contribute equally to the final representation. These differences can arise from variations in biological relevance to the task, data quality (e.g., noise levels or sparsity), or the strength and consistency of the signal. For example, one omics layer may contain highly informative patterns that are strongly correlated with the target phenotype, while the other may be more heterogeneous or affected by experimental noise, potentially diluting the predictive signal.

To account for these disparities, we introduce a mechanism that balances the contributions of intra-omics and interomics information when forming the final node representation. Specifically, we use a weighted integration approach to combine the outputs of the intra-omics and inter-omics GCN modules, as shown in Fig.1(a). This weighted combination allows the model to regulate the influence of each module on the final representation, effectively controlling the flow of information between within-omics and cross-omics signals.

To implement this, we take the hidden features learned from the intra-omics GCNs (**Z**_*u*_ and **Z**_*v*_) and the inter-omics GCNs (**Z**_*v*→*u*_ and **Z**_*u*→*v*_), and integrate them using a weighted sum, defined as:

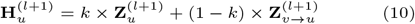

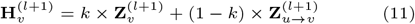

where *k* ∈ (0, 1) is a tunable hyperparameter that controls the relative contributions of intra-omics and inter-omics hidden features. The resulting combined feature representations, 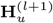 and 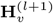, are then used as input for the next layer of SynOmics.

At the final layer, we concatenate the hidden representations and apply a fully connected layer for prediction, as illustrated in Fig.1(c). This operation is defined as:

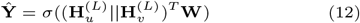

where **Ŷ** ∈ ℝ^*n×nc*^ denotes the predictions, *σ*(.) is the nonlinear activation function, *L* indicates the last layer, and **W** ∈ ℝ^(*p*+*q*)*×nc*^ is the associated weight matrix, with *nc* represents the number of classes. We use the sigmoid activation function for binary classification and the softmax activation function for multi-class classification. The model is trained using binary cross-entropy (*BCE*) loss for binary tasks and categorical crossentropy (*CCE*) loss for multi-class classification:

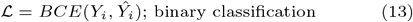

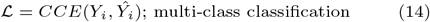

Here, *Y*_*i*_ denotes the ground truth label, and *Ŷ*_*i*_ represents the predicted output for the *i*^*th*^ sample.

This approach enables end-to-end training, where the entire model—including the GCN layers and the fully connected layer—is optimized jointly to generate the final predictions.

### Dual Alignment

The hidden representations generated by the intra-omics and inter-omics modules correspond to the same omics type but are learned through distinct pathways. The intra-omics GCN captures dependencies within a single omics layer, while the inter-omics GCN gathers complementary information from a different omics layer via the bipartite network. Although both modules aim to represent the same set of omics features, they encode different aspects of the data. To ensure consistency between these perspectives, we introduce an alignment loss that minimizes the discrepancy between their outputs, encouraging convergence in a shared feature space. Moreover, to preserve the integrity of the original omics data during transformation, we incorporate a reconstruction loss, which penalizes deviations between the learned intra-omics hidden features and the original inputs. This helps maintain biological interpretability and prevents excessive distortion during feature learning. Together, these loss functions promote coherent integration while preserving meaningful structure in the omics data (Fig.1(b)). To implement this, we initially train the intra-omics and inter-omics GCN modules independently in an unsupervised manner, without using ground truths from the downstream prediction task. This decoupling allows each module to focus solely on learning structural and relational patterns in the data. By separating representation learning from task-specific supervision, we ensure that the learned features retain generalizable biological signals. The total loss function for this pretraining phase is defined as:

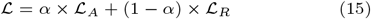

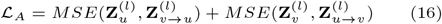

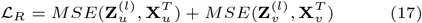

where ℒ_*A*_ and ℒ_*R*_ denote alignment loss and reconstruction loss respectively. The hyperparameter *α* ∈ (0, 1) controls the trade-off between the two losses, and *MSE* refers the mean squared error. After pretraining the GCN modules, we proceed to train SynOmics in an end-to-end manner, where the predictive layer is same as equation (12). The model is then optimized using the loss functions defined in equations (13) and (14) for binary and multi-class classification tasks, respectively.

### Extending to *M* Omics Types

SynOmics is designed to be flexible, supporting the integration of any number of omics data types rather than being limited to just two. To achieve this, we construct an intra-omics network for each of the *M* omics types, along with bipartite networks for every pairwise combination of omics types. For each omics type, the corresponding feature matrix and intra-omics network are processed by an omics-specific GCN to learn intra-omics representations. In parallel, for every omics pair, the two associated feature matrices and their bipartite network are passed through a bipartite GCN to capture cross-omics interactions. After one layer of processing, each omics type yields one hidden representation capturing intra-omics dependencies and *M* − 1 hidden representations that capture its relationships with the other *M* − 1 omics types. These *M* representations are then aggregated using assigned weights and passed to the next layer. In the final layer, the hidden representations from all *M* omics types are combined and passed through a fully connected layer to generate the final prediction.

### Network Construction

We construct the intra-omics adjacency matrices **A**_*u*_ and **A**_*v*_ by computing the pairwise cosine similarity between nodes. Edges are retained only if the cosine similarity exceeds a predefined threshold *ϵ*. Specifically, the adjacency between node *v*_*i*_ and *v*_*j*_ is defined as:

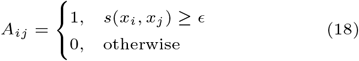

where *x*_*i*_ and *x*_*j*_ are the feature vectors of nodes *v*_*i*_ and *v*_*j*_, respective, and 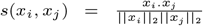 denotes their cosine similarity. The adjacency matrices are then normalized according to equation (2).

We incorporate cross-omics interaction networks obtained from established biological databases as prior knowledge to guide the training of our inter-omics modules. These networks provide valuable biological context, including known molecular interactions, regulatory relationships, and functional associations between different omics features, thereby enhancing the effectiveness of the integration process. In cases where no prior cross-omics interaction data is available, we construct the cross-omics incidence matrices **B**_*u*_ and **B**_*v*_ by computing the pairwise Pearson correlation coefficient (PCC) between features from different omics layers. Edges with a PCC greater than a predefined threshold *ϵ*^*′*^ are retained. Specifically, the adjacency between node *u*_*i*_ from one omics type (e.g., DNA methylation) and node *v*_*j*_ from another omics type (e.g., miRNA) is defined as:

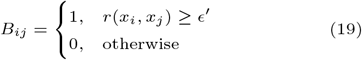

where *x*_*i*_ and *x*_*j*_ are the feature vectors of nodes *u*_*i*_ and *v*_*j*_, respectively, and 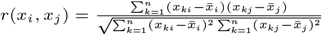 is the PCC between them. Note that for undirected graphs, 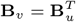. We then construct the bipartite adjacency matrix as defined in equation (3) and normalize it following equation (2).

### Evaluation Methods

#### Disease Classification

To evaluate the binary classification performance of SynOmics, we use two key metrics: **AUROC** (Area Under the Receiver Operating Characteristic Curve) and **MCC** (Matthews Correlation Coefficient).

AUROC provides a comprehensive measure of the model’s ability to distinguish between positive and negative classes across all possible classification thresholds. It evaluates performance based on predicted probabilities, making it particularly informative prior to applying any threshold. This is especially valuable for imbalanced datasets, as it reflects the model’s overall capacity to separate classes probabilistically.

In contrast, MCC evaluates performance after a classification threshold has been applied to convert probabilities into discrete class labels. It considers all components of the confusion matrix, offering a balanced and interpretable summary of prediction quality. This makes MCC particularly effective for assessing final classification outcomes in imbalanced settings.

Since our datasets consist of real-world samples that often exhibit imbalanced class distributions, we selected AUROC and MCC as the most suitable evaluation metrics. To determine the optimal classification threshold, we use Youden’s Index [33], derived from the ROC curve. This method maximizes the sum of sensitivity and specificity, providing a balanced approach to classification performance. Once the optimal threshold is identified, we apply it to the model’s probability predictions to obtain class labels, which are then used to compute the MCC.

#### Survival Analysis

We train a Cox proportional hazards model with an Elastic Net penalty [34] to assess the association between patients’ overall survival and their omics profiles. The Elastic Net penalty combines the *L*_1_-norm and *L*_2_-norm penalties in a weighted manner by maximizing the following log-likelihood function:

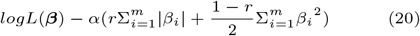

where *L*(***β***) is the partial likelihood of the Cox model, *α* ≥ 0 is a regularization parameter controlling overall shrinkage, and *r* ∈ [0, 1] determines the relative contributions of the *L*_1_ and *L*_2_ penalties. The coefficient *β*_*i*_ corresponds to the *i*-th genomic feature among the *m* features in the omics data. To evaluate model performance, we define high-risk and low-risk groups based on the prognostic index (*PI*) computed from the independent test set. The *PI* represents the linear component of the Cox model, calculated as *PI* = ***β***^*T*^ **X**_*test*_, where **X**_*test*_ is the test set omics profile, and ***β*** is the risk coefficient vector estimated from the model fitted on the training set. Survival outcomes are visualized using Kaplan-Meier survival plots [35]. To construct the high-risk and low-risk groups for these plots, we divide the ordered *PI* values from the test set such that each group contains an equal number of samples. We then use the log-rank test to compare the survival distributions of the two groups and assess whether the observed difference in overall survival is statistically significant.

## Experiments

This section introduces the datasets and networks used in our study, followed by a comparative evaluation of SynOmics against existing integration models. We then assess its performance in survival prediction and biomarker identification. Finally, we analyze the impact of a key hyperparameter and conduct an ablation study to evaluate the individual contributions of intra- and inter-omics learning.

### Datasets and Networks

We applied SynOmics to three TCGA datasets: breast invasive carcinoma (BRCA) [36], lung adenocarcinoma (LUAD) [37], and ovarian serous cystadenocarcinoma (OV) [38]. RNA-seq mRNA expression and miRNA expression data were obtained from the UCSC Xena Hub [39]. DNA methylation data were also collected from the same source to enable experiments involving three omics types. For mRNA expression, we used *log*_2_(*x* + 1) transformed RSEM normalized count data. For miRNA expression, we used *log*_2_(*x* + 1) transformed RPM values. DNA methylation profiles, generated using the Illumina Infinium HumanMethylation450 platform, consisted of beta values representing the ratio of methylated to total probe intensity at each locus. Clinical data for all three cancer types were retrieved from cBioPortal [40]. We obtained the mRNA-miRNA interaction network from TargetScanHuman [41], which provides context++ scores to quantify regulatory relationships between miRNAs and their gene targets.

In the BRCA dataset, patients were classified based on receptor status (Table 2), including estrogen receptor (ER+ vs. ER-), human epidermal growth factor receptor 2 (HER2+ vs. HER2-), progesterone receptor (PR+ vs. PR-), and triplenegative status (TN vs. non-TN). Triple-negative breast cancer patients test negative for all three standard receptors: ER, PR, and HER2. For the LUAD and OV datasets, patients were categorized into short- and long-survival groups based on overall survival time. Patients who survived less than a predefined threshold were placed in the short-survival group, while those who survived longer were classified into the long-survival group. A similar thresholding method was applied to stratify patients into short- and long-duration groups based on disease-free survival. Patients who experienced recurrence before the threshold were assigned to the short-duration group, while those with recurrence after the threshold or no recurrence at all were classified into the long-duration group. Thresholds were selected to ensure that each group contained at least 20 samples. Table 2 summarizes the thresholds used for the LUAD and OV datasets.

**Table 2.**
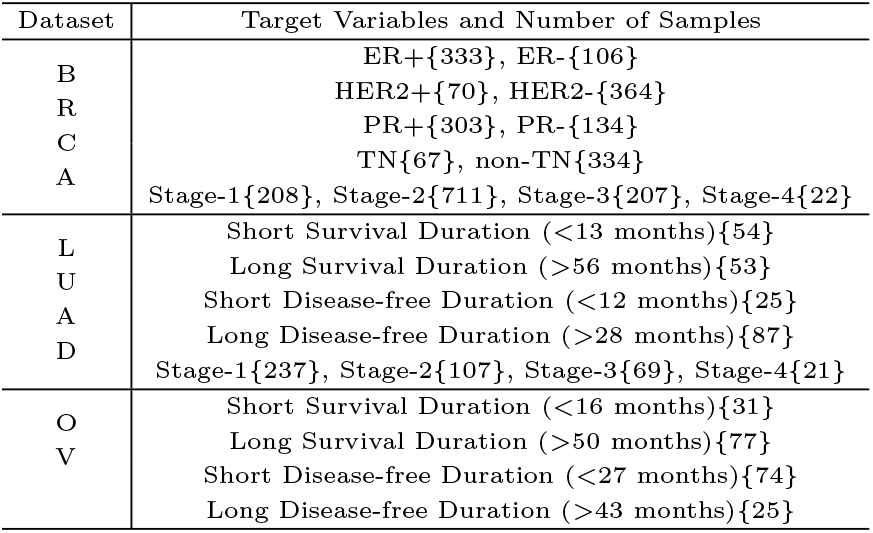
Datasets, target variables, and sample sizes used in this study. The number of samples for each target variable is indicated in curly braces. For the LUAD and OV datasets, the thresholds used to define survival duration groups are shown in parentheses.

In addition to binary classification tasks, we evaluated our model on multi-class classification using cancer stage data from the BRCA and LUAD cohorts. Each dataset includes four stages, representing progressively advanced levels of disease. Table 2 also reports the number of samples associated with each target variable.

The miRNA expression and DNA methylation datasets contained missing values, which we addressed using K-nearest neighbor (KNN) imputation with *K* = 5. Omics datasets typically include thousands of features, many of which exhibit low variability and offer limited analytical value. To reduce noise and improve model performance, we retained only the most variable features, as these are more likely to capture biologically meaningful patterns. Specifically, we selected the top 1,000 most variable features for mRNA expression, the top 200 for miRNA expression, and the top 1,000 for DNA methylation. To further ensure biological relevance, we retained only features with known interactions based on the provided biological networks. We then split the dataset by allocating 80% for training and 20% for testing. The training set was further divided, reserving 20% for validation and using the remaining 80% for model training. After splitting, we standardized the training, validation, and test sets independently to avoid data leakage, scaling each set based on its own Z-scores. This preprocessing results in the input feature matrix *X*_*m*_ ∈ ℝ^*n×d*^, where *n* is the number of samples, *d* is the number of features, and *m* ∈ {1, 2, …, *M}* denotes the omics type. To ensure robust evaluation, we repeated the data split 100 times and reported the average performance metrics across these runs. This procedure helps ensure that our results reflect overall model performance rather than being influenced by random variation in a single split.

### Cancer Outcome Prediction

We compared the classification performance of SynOmics with four existing state-of-the-art multi-omics integration deep learning models: MOGONET [19], MoGCN [17], SUPREME [20], and MOGLAM [21]. All these models utilize GNN architectures that focus on multi-omics integration for various downstream tasks. The classification results for the BRCA, and LUAD and OV datasets are shown in Tables 3 and 4, respectively.

**Table 3.**
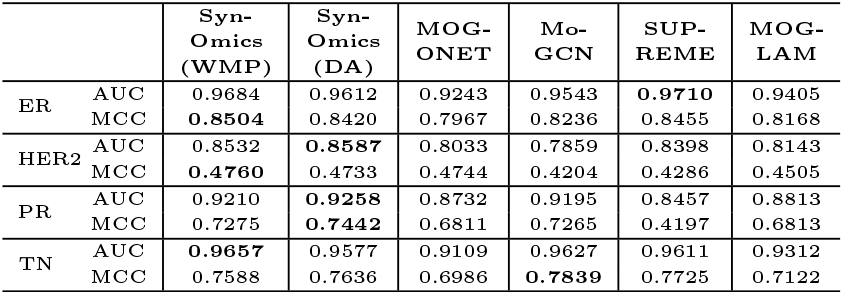
Classification performance on the BRCA dataset for four receptor statuses (ER, HER2, PR, TN). Performance is evaluated using AUC and MCC metrics. ‘WMP’ and ‘DA’ refer to models trained with the weighted message passing and dual alignment strategies, respectively. The best results for each metric are shown in bold.

**Table 4.**
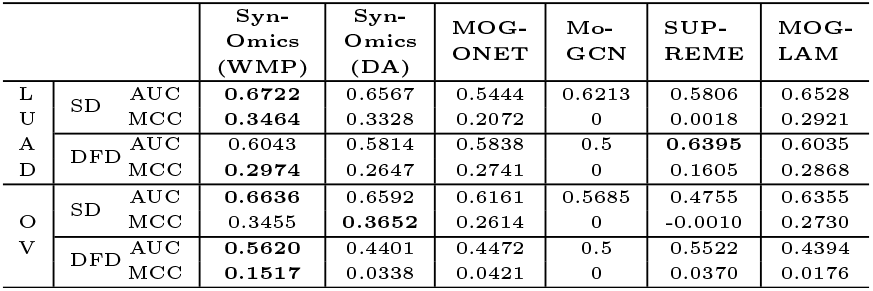
Classification performance on the LUAD and OV datasets for two clinical outcomes: SD and DFD. Performance is evaluated using AUC and MCC metrics. ‘WMP’ and ‘DA’ refer to models trained with the weighted message passing and dual alignment strategies, respectively. The best results for each metric are shown in bold. SD: Survival Duration; DFD: Disease-Free Duration.

As shown in Table 3, SynOmics variants generally achieve better performance than other models on the BRCA dataset. However, for ER status prediction, SUPREME achieves a slightly higher AUC, likely due to its use of sample similarity networks that capture global ranking patterns. This suggests that the ER dataset may contain strong sample-level structure, which benefits models that emphasize overall ranking of cases. In contrast, SynOmics attains a higher MCC, reflecting more accurate binary classification. This may be due to its ability to extract fine-grained, feature-level signals across omics layers. Additionally, SynOmics applies thresholding strategies such as Youden’s Index, which could further contribute to its strong performance in discrete decision-making tasks. For TN status prediction, MoGCN slightly outperforms SynOmics in terms of MCC. This difference may stem from architectural distinctions—MoGCN builds sample similarity networks to guide learning, potentially capturing global relationships among samples. In contrast, SynOmics models feature-level intra- and inter-omics interactions through GCNs and bipartite GCNs, emphasizing regulatory dependencies across omics layers. While SynOmics focuses on fine-grained feature interactions, MoGCN’s sample-level perspective may offer an advantage in capturing broader class separation patterns relevant for TN classification. In the majority of cases, SynOmics outperforms competing models. Notably, its two variants achieve comparable performance, with minor differences, and often lead the benchmarks.

In the LUAD and OV datasets (Table 4), SynOmics demonstrates strong overall performance. SUPREME slightly leads in AUC for LUAD disease-free duration, but both SynOmics variants outperform all other models in the remaining settings. MoGCN’s predictions collapse into a single class, resulting in an MCC of 0.

We also evaluate SynOmics on a multi-class classification task using BRCA and LUAD cancer stage information, as shown in Table 5. Multi-class classification is generally more challenging than binary classification, and thus we observe a performance drop across all models compared to their binary classification results. However, SynOmics with dual alignment outperforms all other methods in BRCA stage prediction, achieving higher AUC and MCC scores. For LUAD stage prediction, MoGCN slightly outperforms SynOmics in terms of AUC, but SynOmics still achieves better performance when considering MCC.

**Table 5.**
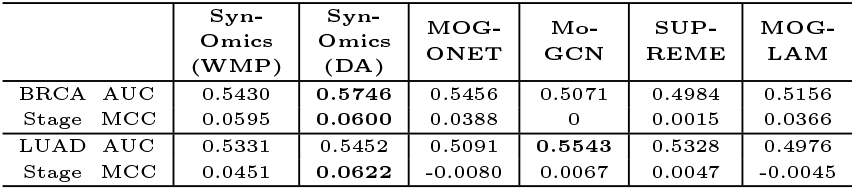
Classification performance on BRCA and LUAD cancer stage groups. Performance is evaluated using AUC and MCC metrics. ‘WMP’ and ‘DA’ refer to models trained with the weighted message passing and dual alignment strategies, respectively. The best results for each metric are shown in bold.

**Table 6.** Ablation results on the BRCA, LUAD, and OV datasets. Performance is evaluated using AUC and MCC metrics. SynOmics is tested in three modes: Both (trained with both intra- and inter-omics modules), Intra (only intra-omics module), and Inter (only inter-omics module). The best results for each metric are shown in bold. SD: Survival Duration; DFD: Disease-Free Duration.

|  |  | AUC |  |  | MCC |  |  |
| --- | --- | --- | --- | --- | --- | --- | --- |
|  |  | Both | Intra | Inter | Both | Intra | Inter |
| BRCA | ER | **0.9684** | 0.9612 | 0.9582 | **0.8504** | 0.8388 | 0.7791 |
|  | HER2 | **0.8532** | 0.8440 | 0.5993 | **0.4760** | 0.4542 | 0.1955 |
|  | PR | **0.9210** | 0.9156 | 0.6995 | **0.7275** | 0.7246 | 0.3815 |
|  | TN | **0.9658** | 0.9594 | 0.8377 | 0.7588 | **0.7638** | 0.4912 |
| LUAD | SD | **0.6722** | 0.6686 | 0.5734 | **0.3464** | 0.3379 | 0.2378 |
|  | DFD | **0.6043** | 0.5332 | 0.5732 | **0.2975** | 0.2272 | 0.2737 |
| OV | SD | **0.6636** | 0.6247 | 0.5411 | **0.3455** | 0.2834 | 0.2498 |
|  | DFD | **0.5620** | 0.4451 | 0.4485 | **0.1517** | 0.0313 | 0.0295 |

### Survival Prediction

We conducted an overall survival prediction analysis using the BRCA, LUAD, and OV datasets to assess the quality of the hidden representations generated by SynOmics. For this analysis, we extracted the hidden representations produced by SynOmics before the predictive layer and input them into the

Cox proportional hazards model, as outlined in Section *Survival Analysis*. We set the relative weight *r* in equation (20) to 0.5 to combine the subset selection property of the *L*_1_-norm with the regularization strength of the *L*_2_-norm. The Kaplan–Meier survival plots and log-rank test *p*-values in Fig.2 demonstrate that the hidden representations generated by SynOmics possess strong prognostic ability in distinguishing between high-risk and low-risk groups. For example, at approximately 5 years (60 months), the high-risk group in the BRCA dataset exhibited a survival probability of 69.84% (95% CI: 47.78%–83.99%), whereas the low-risk group showed a higher survival probability of 90.91% (95% CI: 50.81%–98.67%). In the LUAD dataset, the high-risk group demonstrated a markedly lower survival probability of 9.09% (95% CI: 0.54%–33.29%) compared to 81.82% (95% CI: 44.74%–95.12%) in the low-risk group. Similarly, for the OV dataset, the high-risk group had a survival probability of 27.27% (95% CI: 6.52%–53.89%), while the lowrisk group maintained a survival probability of 80.81% (95% CI: 42.35%–94.85%). All of these separations were statistically significant (*p*-value *<* 0.05). As indicated by the *p*-values, none of the datasets could significantly distinguish between high-risk and low-risk groups in their original form (see Supplementary Section S1). In contrast, the hidden representations generated by SynOmics demonstrate improved prognostic capability. They enable a significant separation between the two risk groups, highlighting their effectiveness for downstream survival analysis tasks.

**Fig 2.**
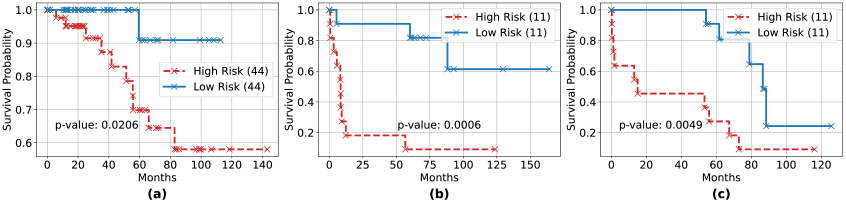
Survival analysis based on the learned hidden representations for (a) BRCA (ER status), (b) LUAD (survival duration), and (c) OV (survival duration) datasets. The numbers in parentheses indicate the sample sizes for the low- and high-risk groups. The p-value is computed using the log-rank test to assess the difference in overall survival between the two groups. *Alt Text: Kaplan–Meier curves generated using SynOmics representations for stratifying patient risk across BRCA, LUAD, and OV datasets, with group sizes noted and log-rank p-values shown*.

### Biomarker Identification

SynOmics begins with individual omics types and combines intra- and inter-layer signals to learn a unified hidden representation that captures complex relationships between omics features. This representation not only supports disease classification but also facilitates the identification of discriminative biomarkers. To evaluate its effectiveness, we conducted a feature-level analysis to examine how well the hidden representation distinguishes between different lung cancer (LUAD) stages.

Our input data consisted of 1,200 features—1,000 from mRNA expression and 200 from miRNA expression. Throughout the SynOmics layers, we maintained the same feature-space dimension to maximize the benefits of information sharing through the feature networks. As a result, the output hidden representation retained the same number of features, allowing for a direct comparison between the input and output features of SynOmics. To identify the most predictive features, we conducted ANOVA tests on both the 1,200-dimensional input and output feature sets to determine which features, in both the original and encoded spaces, are significantly associated with cancer stage. Specifically, we grouped the input and output feature expressions based on their corresponding stages and conducted ANOVA tests to determine which features exhibited the most statistically significant (*p*-value *<* 0.05) differences in expression. This analysis revealed 8 significant features in the original dataset and 33 significant features in the hidden representation, demonstrating that SynOmics enhances omics data representation by uncovering additional features strongly associated with cancer progression. For further analysis, we categorized patients into high- and low-expression groups based on each significant feature identified by SynOmics and conducted log-rank tests to evaluate their ability to distinguish survival differences between these groups.

Fig.3(a) provides a violin plot illustrating the expression pattern of one representative significant feature (among the 33 identified by SynOmics) across advancing cancer stages, while Fig.3(b) shows patient survival probabilities for high- and low-expression groups based on the same significant feature. This feature exhibited a statistically significant variation in expression levels across cancer stages, as determined by the ANOVA test (*p*-value = 0.0183). The mean expression values showed a progressive increase across stages, with stage 1 having a mean of 0.2701 (95% CI: 0.1368–0.4035), stage 2 with 0.6138 (95% CI: 0.1857–1.0419), stage 3 with 0.6400 (95% CI: 0.0868–1.1931), and stage 4 reaching 1.2625 (95% CI: 0.0903–2.4347). This pattern suggests a potential upregulation of the feature with disease progression. Survival analysis further demonstrated the prognostic utility of the representative feature. At approximately 5 years (60 months), the high-risk group showed a survival probability of 32.94% (95% CI: 16.72%–50.18%), whereas the low-risk group showed a survival probability of 59.46% (95% CI: 20.67%–84.25%). The log-rank test confirmed that this separation was statistically significant (*p*-value = 0.0037). Together, these results indicate that the representative feature may serve as both a stage-associated and prognostically relevant biomarker, with robust evidence from both expression variability and survival stratification.

**Fig 3.**
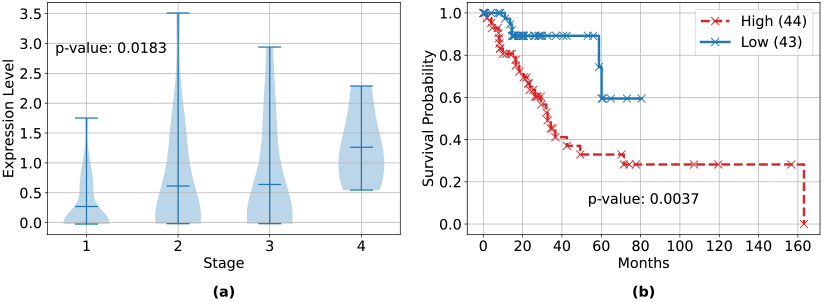
Representative results from biomarker identification. (a) Violin plot showing the expression levels of a SynOmics-selected feature across progressively advancing cancer stages. The p-value is computed using the ANOVA test. (b) Kaplan-Meier survival plot comparing overall survival between patient groups with high and low expression of the same feature. Sample sizes for each group are shown in parentheses. The p-value is calculated using the log-rank test. *Alt Text: A two-part figure displaying statistical validation of a SynOmics-selected biomarker: (a) expression variation across cancer stages via violin plot and (b) survival difference via Kaplan–Meier curves for expression-defined groups*.

### Hyperparameter Tuning

In the ‘weighted message passing’ module of SynOmics, *k* determines the relative contribution of intra-omics and interomics features when combining the two sources of information, as described in equations (10) and (11). If the value of *k* is too large, the hidden representation tends to reflect mostly the intra-omics dynamics, with minimal influence from the other omics types. Conversely, if *k* is too small, the hidden representation predominantly captures information from other omics sources, neglecting the intra-omics dynamics. To effectively capture both types of dynamics, a balance between the two sources of information is essential. Fig.4 shows the model’s performance (AUC and MCC scores) across different values of *k* for the three datasets. It is evident that lower values of *k* lead to poorer performance, while performance improves as *k* increases. The best results are typically observed when *k* is in the range of 0.4 to 0.5. However, when *k* becomes too large, performance declines again, particularly in the LUAD and OV datasets. This suggests that neither intra-omics nor inter-omics dynamics alone provides optimal performance; rather, combining both sources leads to the best results.

**Fig 4.**
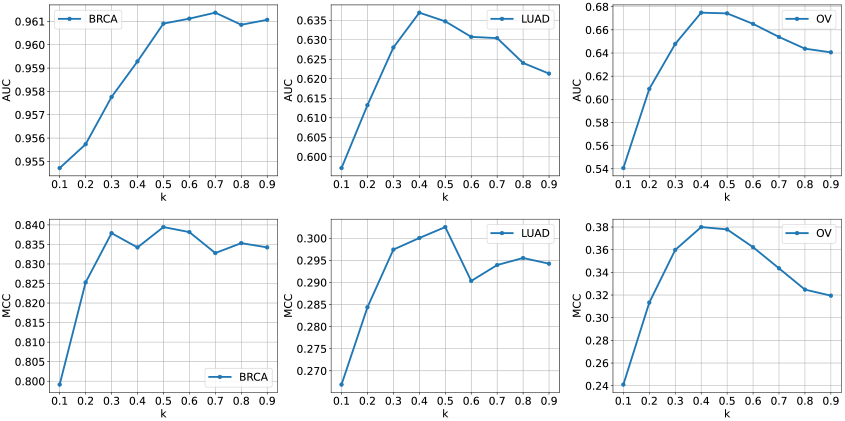
SynOmics performance under varying intra-omics and inter-omics weights. AUC (top row) and MCC (bottom row) scores for the BRCA (ER status), LUAD (survival duration), and OV (survival duration) datasets across different values of k, which controls the relative contribution of intra-omics and inter-omics representations. *Alt Text: Figure showing how SynOmics’ classification and survival prediction performance changes as the balance between intra- and inter-omics information is adjusted, using AUC and MCC as metrics*.

The analysis of the hyperparameter *α* in the ‘dual alignment’ module, along with other hyperparameter settings, is available in Supplementary Section S2.

### Ablation Study

To assess the relative importance of the intra-omics and interomics modules in SynOmics, we conduct an ablation study to better understand the contributions of each component. In this study, we train SynOmics using only the intra-omics or interomics modules individually.

For intra-omics learning, we focus exclusively on constructing omics-specific networks and applying the corresponding GCNs, without incorporating any cross-omics interactions through the bipartite network. In this configuration, only the GCNs defined in equations (6) and (7) are trained, while the bipartite GCNs in equations (8) and (9) are excluded. As a result, the model relies entirely on intra-omics representations, with no involvement of weighted message passing or the dual alignment module.

In contrast, for inter-omics learning, we rely solely on the bipartite interaction networks and their associated GCNs to model cross-omics relationships. In this case, omics-specific networks are not used, and only the bipartite GCNs in equations (8) and (9) are trained. As with the intra-omics setup, the weighted message passing and dual alignment modules are omitted, since integration across omics types is not performed. The outcomes of this ablation study are summarized in Table

6. The table clearly shows that the model trained with both modules outperforms the versions trained with just one module. When trained to capture only intra-omics relationships, the model experiences a noticeable performance drop. However, the drop is even more significant when the model is trained solely on inter-omics relationships. This suggests that focusing only on inter-omics dynamics, without considering intra-omics interactions, is insufficient for effective multi-omics integration. While considering only intra-omics relationships may yield decent results, the most effective multi-omics integration requires accounting for both intra- and inter-omics dynamics.

### Results for Three Omics Types

To demonstrate the extensibility of SynOmics, we expanded it to incorporate three omics types. In addition to the mRNA and miRNA expression data, we included DNA methylation data as the third omics type. Including more omics types often reduces the usable sample size in supervised learning, as complete data across all sources is required—a constraint that affected the BRCA dataset. For LUAD, however, the DNA methylation samples aligned with the existing set due to prior thresholding. Thus, we used LUAD for the three-omics analysis to ensure a fair comparison with the two-omics results. It is important to note that all the other models being compared were also trained with the three types of omics data.

To build the mRNA expression-DNA methylation and miRNA expression-DNA methylation bipartite networks, we applied the method outlined in equation (19). The experimental results are presented in Table 7. It is evident that SynOmics outperforms the other models when trained with the additional DNA methylation data. Furthermore, as shown in Table 4 and Table 7, the inclusion of extra omics data improved the performance of SynOmics.

**Table 7.**
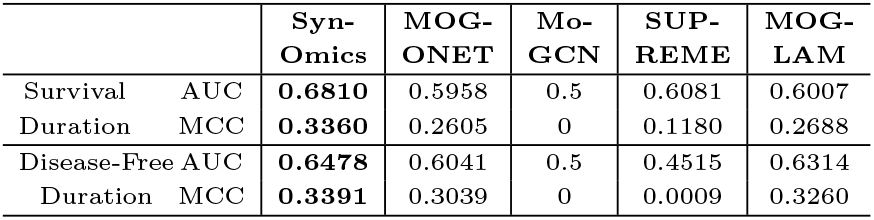
Classification performance on the LUAD dataset using three omics types. Results are evaluated using AUC and MCC metrics. The best results for each metric are shown in bold.

## Discussion

Multi-omics integration is essential for uncovering coordinated biological mechanisms across layers such as genomics, transcriptomics, proteomics, and metabolomics. Traditional methods that rely on sample correlations often fail to capture the true feature-level relationships that drive biological processes. Sample correlation, while useful in some contexts, does not necessarily reflect the true dynamics of feature-level interactions within biological systems. Sample networks are typically built on the assumption that there exists homogeneity within samples and consistent relationships across varying biological states. This assumption can be erroneous, especially in low-sample, high-dimensional contexts typical of omics data, where sample variability may overshadow true biological signals.

Rather than depending on correlations between samples, SynOmics emphasizes feature-level interactions, improving its resilience to changes in sample size and variability. It achieves cross-omics integration through bipartite networks, which represent relationships between features across omics types and can be built using known interactions or computed similarities. This approach is better equipped to incorporate contextual information—such as the regulatory role of miRNA in mRNA expression—compared to context-agnostic methods that use fully connected layers or autoencoders. Experimental results demonstrate the superiority and effectiveness of SynOmics. Notably, its survival prediction capabilities indicate its potential for patient survival analysis. Effective multi-omics integration should consider both intra-omics and inter-omics interactions, as evidenced by the ablation studies conducted with SynOmics. The model adeptly captures how expression levels correlate with different disease stages.

SynOmics operates at the feature level, so its computational complexity is largely determined by the feature size of the omics data used. Specifically, the complexity is mathematically expressed as *O*(*lp*^2^*n*), where *l* represents the number of layers, *p* represents the feature size of the omics type with the larger dimension, and *n* represents the sample size. The quadratic nature of this complexity with respect to *p* can be significant, especially when dealing with large feature sets. However, applying dimensionality reduction techniques can help mitigate this issue by reducing the number of features. In many omics datasets, a large portion of features have low variance and contribute minimally to key tasks such as disease or survival prediction. In this study, we therefore focused on using features with high variance to train the model. We also incorporated early stopping in our model training based on validation loss. If the validation loss does not improve over a specified number of epochs (e.g., 20 epochs), we consider the loss to be saturated and terminate the training. This approach also helps reduce training time. As a result, our model achieved the shortest training time compared to the others. The average training times for SynOmics, MOGONET, MoGCN, SUPREME, and MOGLAM were 5.35, 131.10, 15.30, 174.32, and 199.37 seconds, respectively, measured on a machine with an AMD Ryzen 9 5950X CPU, 128 GB RAM, and a single NVIDIA RTX A4500 (20 GB) GPU.

Parallel integration of information across all omics layers—rather than at the beginning or end of the process—enables SynOmics to create a cohesive, integrated representation. To further improve the performance of SynOmics, it may be advantageous to explore advanced network creation techniques [42, 43, 44] beyond traditional methods such as cosine similarity or correlation, and to implement more sophisticated imputation strategies [45, 46, 47, 48, 49] instead of relying solely on KNN. Establishing an effective bipartite network between omics types presents challenges, but when accomplished, it can greatly improve model performance by accurately reflecting the multilayered interactions among diverse biological features. In future work, we plan to extend SynOmics for application in single-cell and spatial multi-omics, where spatial coordinates and cell-level similarities can be incorporated through appropriate network modeling.

In conclusion, our findings highlight that SynOmics is a flexible and effective model for multi-omics integration, enabling a more in-depth understanding of complex diseases. By leveraging the dynamic relationships within omics data, the model enhances the accuracy of disease prediction and uncovers deeper insights into how different molecular factors contribute to disease progression. SynOmics can identify key biomarkers, which are essential for advancing medical research. These biomarkers have the potential to guide the development of targeted therapies, leading to more effective drugs and treatments. Overall, SynOmics provides an effective and scalable approach to multi-omics integration, offering a path toward more accurate biological interpretation and clinically actionable outcomes.

## Supporting information

Supplementary Section S1, Supplementary Section S2

**Key Points**

- SynOmics integrates multi-omics data at the feature level using both intra-omics networks and inter-omics bipartite networks.
- SynOmics incorporates two alternative integration approaches and adopts a parallel learning strategy for joint representation learning.
- SynOmics consistently outperforms state-of-the-art methods in classification tasks and demonstrates robust performance in survival prediction.
- SynOmics identifies biologically relevant features associated with disease progression and differential expression across stages.

## Competing interests

No competing interest is declared.

## Author contributions statement

M.N.A. and W.Z. designed the SynOmics model. M.N.A. implemented the algorithm and drafted the manuscript. W.Z. supervised the work and provided suggestions for improving both the model and the manuscript.

## Acknowledgments

The results are based upon data generated by The Cancer Genome Atlas established by the NCI and NHGRI. Information about TCGA and the investigators and institutions who constitute the TCGA research network can be found at http://cancergenome.nih.gov. The dbGaP accession number to the specific version of the TCGA dataset is phs000178.v11.p8. This work was supported by grants from the National

Science Foundation (NSF) [NSF-III2246796 and NSF-III2152030]

**Muhtasim Noor Alif** is a doctoral student in the Department of Computer Science at the University of Central Florida. His research centers on applying machine learning techniques to computational biology.

**Wei Zhang** is an associate professor in Computer Science at the University of Central Florida. He received his PhD in Computer Science from University of Minnesota Twin Cities. His research interests are broadly in cancer genomics, network-based learning in bioinformatics, and disease phenotype prediction.

## Notes

### Competing Interest Statement

The authors have declared no competing interest.

