## Supplementary Section S1, Supplementary Section S2 for "SynOmics: Integrating Multi-omics Data Through Feature Interaction Networks"

### S1 Survival Analysis

Figure S1 illustrates the survival prediction performance derived from the original datasets for breast (BRCA), lung (LUAD), and ovarian (OV) cancers.

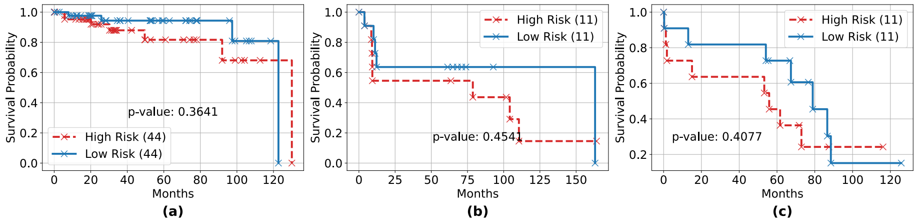


Figure S1: Survival analysis based on original input features for (a) BRCA (ER status), (b) LUAD (survival duration), and (c) OV (survival duration) datasets. The numbers in parentheses indicate the sample sizes for the low- and high-risk groups. The $p$-value is computed using the log-rank test to assess the difference in overall survival between the two groups.

### S2 Hyperparameter Tuning

In the 'dual alignment' module, the hyperparameter $\alpha$ plays a key role during training. When $\alpha$ is too large, the model primarily focuses on aligning the two hidden representations (intra and inter-omics). On the other hand, if $\alpha$ is too small, the model places more importance on ensuring the hidden representation is close to the original data. In our experiments, as shown in Figure S2, the scores with respect to $\alpha$ initially appear unpredictable. However, upon closer inspection, we can see that the variations in both AUC and MCC are minimal. This suggests that the model maintains robust performance across different values of $\alpha$.

We also describe additional hyperparameter settings used for training SynOmics. To identify the optimal configuration, we constructed a parameter grid with multiple candidate values for each hyperparameter and trained the model across 100 random splits. The common hyperparameters shared between both modules of SynOmics, as well as those specific to the dual alignment module, are summarized in Table S1. The hyperparameter combinations for the best 10 models of the 'weighted message passing' module are shown in Table S3. The hyperparameter combinations for the 10 best-performing models of the 'dual alignment' module are shown in Table S2.

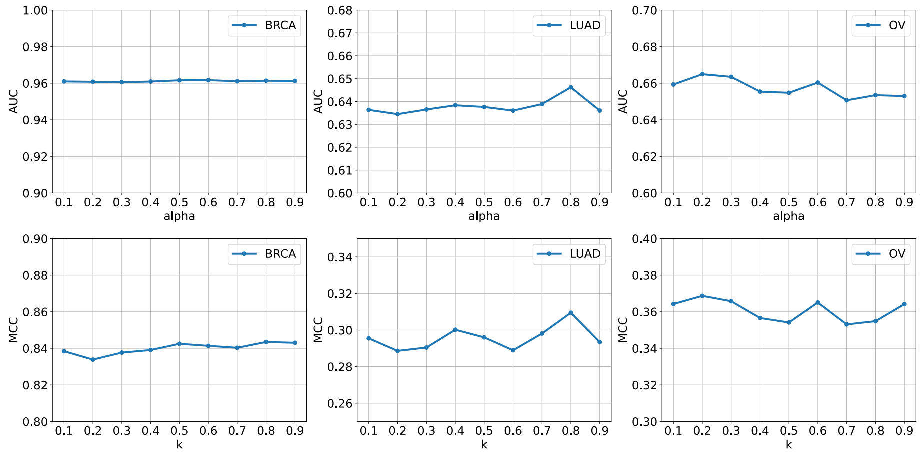


Figure S2: AUC and MCC scores for the BRCA (ER status), LUAD (survival duration), and OV (survival duration) datasets at different values of the ‘dual alignment’ hyperparameter $\alpha$, which controls the balance between alignment and reconstruction loss..

| | **Hyperparameter** | **Description** |
|---|---|---|
| Common Parameters | Number of layers | Number of SynOmics layers used during training. |
| | Batch size | Batch size used for feeding input data during training. |
| | Learning rate | Learning rate used by the optimizer. |
| | Epochs | Total number of training epochs. |
| | Adjacency threshold | Threshold used to binarize the graph adjacency matrix. |
| | Hidden dimension | Dimensionality of the hidden layer in the fully-connected prediction layer. |
| | Bias | Indicates whether the model includes a bias term. |
| WMP Parameters | $k$ | Controls the weight for intra-omics contribution during integration of intra- and inter-omics hidden representations. |
| DA Parameters | Pretraining epochs | Number of epochs used for pretraining the SynOmics layers before alignment. |
| | $\alpha$ | Weight assigned to the alignment loss in the combined loss function (alignment + reconstruction). |

Table S1: Descriptions of the hyperparameters used to train SynOmics. ‘WMP’ refers to weighted message passing, and ‘DA’ refers to dual alignment.

| Number of Layers | Batch Size | Learning Rate | Epochs | Pretraining Epochs | Adjacency Threshold | Hidden Dimension | Bias | $\alpha$ |
|---|---|---|---|---|---|---|---|---|
| 3 | 32 | 0.001 | 50 | 10 | 0.0 | 64 | True | 1.0 |
| 1 | 32 | 0.001 | 10 | 10 | 0.1 | 32 | True | 1.0 |
| 1 | 32 | 0.001 | 50 | 30 | 0.0 | 64 | True | 1.0 |
| 1 | 32 | 0.001 | 50 | 50 | 0.1 | 32 | True | 0.0 |
| 1 | 32 | 0.001 | 30 | 10 | 0.1 | 64 | True | 0.5 |
| 1 | 32 | 0.001 | 50 | 30 | 0.0 | 32 | True | 0.0 |
| 1 | 32 | 0.0001 | 50 | 10 | 0.1 | 32 | True | 0.0 |
| 1 | 32 | 0.001 | 30 | 10 | 0.0 | 32 | True | 0.8 |
| 1 | 32 | 0.0001 | 50 | 10 | 0.0 | 32 | True | 0.8 |
| 1 | 32 | 0.001 | 10 | 30 | 0.0 | 64 | True | 1.0 |

Table S2: Top 10 performing model hyperparameters for the 'dual alignment' module.

| Number of Layers | Batch Size | Learning Rate | Epochs | Adjacency Threshold | Hidden Dimension | Bias | $k$ |
|---|---|---|---|---|---|---|---|
| 1 | 32 | 0.001 | 50 | 0.1 | 64 | True | 0.8 |
| 1 | 32 | 0.0001 | 50 | 0.0 | 64 | True | 0.7 |
| 1 | 32 | 0.001 | 50 | 0.5 | 64 | True | 0.9 |
| 5 | 32 | 0.0001 | 30 | 0.0 | 64 | True | 0.7 |
| 3 | 32 | 0.001 | 100 | 0.0 | 32 | True | 0.6 |
| 1 | 32 | 0.0001 | 100 | 0.1 | 32 | True | 0.5 |
| 2 | 32 | 0.0001 | 100 | 0.5 | 64 | True | 0.9 |
| 2 | 32 | 0.0001 | 30 | 0.5 | 64 | True | 0.9 |
| 1 | 32 | 0.001 | 30 | 0.5 | 64 | True | 0.8 |
| 1 | 32 | 0.001 | 50 | 0.5 | 64 | True | 0.5 |

Table S3: Top 10 performing model hyperparameters for the 'weighted message passing' module.
